# TreeQTL: hierarchical error control for eQTL findings

**DOI:** 10.1101/021170

**Authors:** Christine Peterson, Marina Bogomolov, Yoav Benjamini, Chiara Sabatti

**Affiliations:** Department of Health Research and Policy, Stanford University, Stanford, CA 94305, U.S.A.; Faculty of Industrial Engineering and Management, Technion, Haifa, 32000, Israel.; Department of Statistics and Operations Research, Tel Aviv University, Tel Aviv, 69978, Israel.; Departments of Health Research and Policy and Statistics, Stanford University, Stanford, CA 94305, U.S.A.

## Abstract

**Summary:** Commonly used multiplicity adjustments fail to control the error rate for reported findings in many expression quantitative trait loci (eQTL) studies. TreeQTL implements a stage-wise multiple testing procedure which allows control of appropriate error rates defined relative to a hierarchical grouping of the eQTL hypotheses.

**Availability and Implementation:** The R package TreeQTL is available for download at http://bioinformatics.org/treeqtl.

## Introduction

The overarching goal of eQTL analysis is to gain insight into the genetic regulation of gene expression. Typically, this is carried out by testing a vast collection of hypotheses *H*_*vp*_ probing association (or linkage) between the genotype at variant (locus) *v* and the measured expression for probe *p*, where *v* = 1, …, *M*, *p* = 1, …, *G*, *M* is on the order of hundreds of thousands, and *G* of tens of thousands. Given the large number of hypotheses tested, the need to adjust for multiplicity is universally recognized and the false discovery rate (FDR) [5] is typically adopted as the target global error rate.

In an effort to improve interpretability, reporting of results is typically organized along more general findings such as the discovery of genes subject to local regulation [7] or regulatory SNPs (eSNPs) [9]. The adopted strategy for multiplicity adjustment needs to offer guarantees on the specific reported discoveries. For example, imagine testing the *H*_*vp*_ hypotheses using the Benjamini Hochberg (BH) [5] rule and defining as an eSNP those variants *v* for which *H*_*vp*_ is rejected for at least some probe *p*. While this would control the FDR among the *H*_*vp*_ rejections, it would offer no control of the rate of false eSNP discoveries, as shown by the simulations in [10].

Researchers in the eQTL field have recognized this challenge and have noted that since local regulation is more common than distal [1], hypotheses probing these two mechanisms should be tested separately. However, there is no single standard in the literature for error rates targeted or error-controlling strategies: for example, one finds the notion of per-gene error rates [9, 13] or the application of Bonferroni across genes [14] in local regulation, while for distal effects significance cut-offs vary from 5 × 10^−8^ [8] to 5.78 × 10^−12^ [14]. This makes comparison across studies and replicability challenging.

## 1 Approach

To overcome the confusion generated by the plurality of approaches and to provide guarantees relative to the discoveries reported, it is useful to recognize the structure among the hypotheses tested in an eQTL study and to introduce some terminology. We distinguish between hypotheses testing local (when the distance between variant *v* and probe *p* is less then a threshold) and distal regulation, indicating them with *L*_*vp*_ and *D*_*vp*_, respectively.

Further, we recognize that we might be interested mainly in identifying which genes appear to have local (distal) regulation prior to obtaining a detailed list of the variants involved in this regulation; or we might want to pinpoint SNPs that appear to have local (distal) effects on multiple genes, even without committing to a comprehensive list of these genes. Recalling that our measurements of expression are mediated by probes and that typically typed genetic variants are SNPs, we use LeProbe to signify a probe whose expression is influenced by some local DNA variants and LeSNP to designate a specific SNP associated to variability in expression for some local probes. With LeAssociation we signify the local association between one specific SNP and one specific probe. For distal regulation, DeProbe, DeSNP, and DeAssociation are similarly defined. Mapping these back to the original collection of hypotheses, we note, for example, that to discover a DeSNP is equivalent to rejecting the intersection hypothesis *D*_*v*•_=∩_*p*_ D_*vp*_.

To control global error rates defined in terms of the reported discoveries (LeSNPs, LeAssociations, etc.), we have implemented in TreeQTL a multi-resolution approach based on results in [3] and whose practical effectiveness in GWAS has been described in [10]. Furthermore, by accounting for the described structure, TreeQTL has the potential to increase power: the effective number of tests is reduced and one can capitalize on the adaptivity of FDR.

## 2 Methods

TreeQTL is a hierarchical testing procedure that distinguishes two levels of discoveries within each class of local or distal regulation. In Level 1, the users can specify their primary interest as the identification of either eProbes or eSNPs. Given this choice, all the pair-wise association hypotheses are given a position in a tree similar to that shown in Figure 1: each eProbe or eSNP hypothesis indexes the collection of simple hypotheses by whose intersection it is defined. Level 1 hypotheses are tested controlling for global errors within the two regulation classes. The more granular eAssociation hypotheses in Level 2 are tested only when they correspond to a global hypothesis rejected in Level 1.

**Figure 1:**
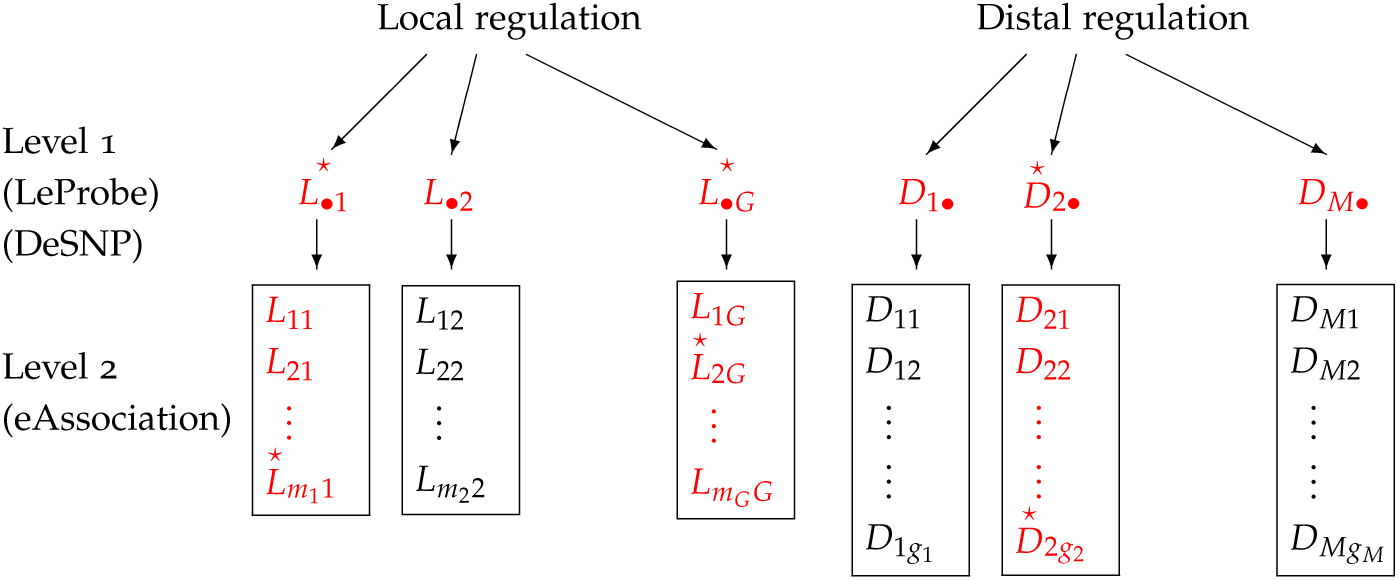
Organization of eQTL hypotheses in TreeQTL. Hypotheses testing local regulation are on the left and hypotheses testing distal regulation are on the right. Local regulation hypotheses have been grouped by probe and distal regulation hypotheses are grouped by SNP, so that Level 1 rejections will result in discoveries of LeProbes and DeSNPs. Tested hypotheses are colored in red, and rejected hypotheses indicated with a star.

TreeQTL takes as input the *p*-values for each hypothesis in Level 2: these may be obtained via Matrix eQTL [11] and their correctness is of crucial importance. The *p*-values for the Level 1 hypotheses are computed using Simes’ rule [12] on the families they index. This summary of the evidence for the global null hypotheses is relatively robust to dependence [4]. Users can, however, input alternative *p*-values for Level 1, such as those obtained via permutation.

Testing proceeds from Level 1, where the default option is to control the FDR at a user-specified level *q*_1_ using the BH [5]. This guarantees control of the FDR for Level 1 discoveries when all hypotheses are independent or under positive regression dependence. The Benjamini-Yekutieli procedure [6], robust to any dependence between tests, is available as an alternative. A third, more stringent option is to control the family-wise error rate (FWER) using the Bonferroni correction.

In Level 2, significance is established using BH within each set of eAssociation hypotheses corresponding to an eSNP or eProbe identified at Level 1, at the more stringent level needed to account for selection [3]. The expected average proportion of false discoveries across the selected Level 2 families is then controlled to the user-specified target level *q*_2_.

## 3 Example application

To demonstrate feasibility, we applied TreeQTL to whole-blood data from the pilot phase of the GTEx project [2]: genotype data at 6,820,472 SNPs and expression levels for 30,115 genes are available across 156 subjects. Following the steps in [2], *p*-values were obtained by applying Matrix eQTL [11] to quantile-normalized gene expression, adjusting for both known and unknown technical covariates (note that power for distal analysis is likely to be limited with this sample size). In applying TreeQTL, we identified eSNPs as the discovery of interest in Level 1, and set *q*_1_ = *q*_2_ = 0.05 for local and *q*_1_ = *q*_2_ = 0.01 for distal regulation. This led to the discovery of 199,032 LeSNPs, involved in 325,529 LeAssociations, and 164,860 DeSNPs involved in 216,933 DeAssociations. By comparison applying BH at levels *q* = 0.05 and *q* = 0.01 across all local and distal hypotheses, respectively, led to the identification of 250,945 LeSNPs (with 389,507 LeAssociations) and 179,625 DeSNPs (with 216,683 DeAssociations). TreeQTL, then, results in a similar number of eAssociations, but these are grouped under a small number of eSNPs.

## 4 Conclusion

By analyzing local and distal regulation separately, TreeQTL has less stringent cut-offs for tests probing local effects, where it is expected that a larger number of hypotheses will be non-null. By grouping hypotheses relative to the same SNP or the same probe, TreeQTL capitalizes on the inherent heterogeneity of the problem. For example, while few SNPs will act as distal regulators of expression and might influence multiple probes, the vast majority of SNPs will not have such an effect: testing all the eAssociation hypotheses relative to one SNP together, separately from those concerning other loci, allows greater power for SNPs with true regulatory roles and reduces false positives. Finally, the hierarchical structure of TreeQTL assures control of the FDR for eSNP and eProbe discoveries. While the current version of TreeQTL implements methodology relative to studies involving only one tissue, future releases will incorporate approaches for the more complex structure of multi-tissue investigations.

## Acknowledgements

The pilot release of the GTEx data (accession no. phs000424.v3.p1) is available through dbGaP (http://www.ncbi.nlm.nih.gov/gap).

### Funding

This work was supported by the National Institutes of Health [MH101782 to C.P. and C.S., HG006695 to C.S. and Y.B.]; and the Israel Science Foundation [1112/14 to M.B.].

## References

[1] F. Albert and L. Kruglyak (2015) The role of regulatory variation in complex traits and disease. Nature Reviews Genetics, 16, 197–212.

[2] K. G. Ardlie, et al. (2015) The Genotype-Tissue Expression (GTEx) pilot analysis: multitissue gene regulation in humans. Science, 8, 648–660.

[3] Y. Benjamini and M. Bogomolov (2014) Selective inference on multiple families of hypotheses. JRSS B, 76, 297–318.

[4] Y. Benjamini and R. Heller (2008) Screening for partial conjunction hypotheses. Biometrics, 64, 1215–1222.

[5] Y. Benjamini and Y. Hochberg (1995) Controlling the false discovery rate: a practical and powerful approach to multiple testing. JRRS B, 57, 289–300.

[6] Y. Benjamini and D. Yekutieli (2001) The control of the false discovery rate in multiple testing under dependency. Ann. Statist., 29, 1165–1188.

[7] H. H. Göring, J. E. Curran, M. P. Johnson, T. D. Dyer, J. Charlesworth, S. A. Cole, J. B. Jowett, Lawrence, J. Abraham, D. L. Rainwater, A. G. Comuzzie, M. C. Mahaney, L. Almasy, W. MacCluer, A. H. Kissebah, G. R. Collier, E. K. Moses and J. Blangero (2007) Discovery of expression QTLs using large-scale transcriptional profiling in human lymphocytes. Nat. Genet., 39: 1208–1216.

[8] E. Grundberg, K. S. Small, A. K. Hedman, A. C. Nica, A. Buil, S. Keildson, J. T. Bell, T. P. Yang, E. Meduri, A. Barrett, J. Nisbett, M. Sekowska, A. Wilk, S. Y. Shin, D. Glass, M. Travers, L. Min, S. Ring, K. Ho, G. Thorleifsson, A. Kong, U. Thorsteindottir, C. Ainali, A. S. Dimas, N. Hassanali, C. Ingle, D. Knowles, M. Krestyaninova, C. E. Lowe, P. Di Meglio, S. B. Montgomery, L. Parts, S. Potter, G. Surdulescu, L. Tsaprouni, S. Tsoka, V. Bataille, R. Durbin, F. O. Nestle, S. O?Rahilly, N. Soranzo, C. M. Lindgren, K. T. Zondervan, K. R. Ahmadi, E. E. Schadt, K. Stefansson, G. D. Smith, M. I. McCarthy, P. Deloukas, E. T. Dermitzakis and T. D. Spector (2012) Mapping *cis*- and *trans*-regulatory effects across multiple tissues in twins. Nat. Genet., 44, 1084–1089.

[9] A. C. Nica, S. B. Montgomery, A. S. Dimas, B. E. Stranger, C. Beazley, I. Barroso, and E. T. Dermitzakis (2010) Candidate causal regulatory effects by integration of expression QTLs with complex trait genetic associations. PLoS Genet., 6, e1000895.

[10] C. Peterson, M. Bogomolov, Y. Benjamini and C. Sabatti (2015) Many phenotypes without many false discoveries: error controlling strategies for multi-trait association studies. arXiv:1504.00701

[11] A. Shabalin (2012) Matrix eQTL: ultra fast eQTL analysis via large matrix operations. Bioinformatics, 28, 1353–1358.

[12] R. Simes (1986) An improved Bonferroni procedure for multiple tests of significance. Biometrika, 73, 751–754.

[13] B. E. Stranger, S. B. Montgomery, A. S. Dimas, L. Parts, O. Stegle, C. E. Ingle, M. Sekowska, G. D. Smith, D. Evans, M. Gutierrez-Arcelus, A. Price, T. Raj, J. Nisbett, A. C. Nica, C. Beazley, R. Durbin, P. Deloukas and E. T. Dermitzakis (2012) Patterns of *cis* regulatory variation in diverse human populations. PLoS Genet., 8, e1002639.

[14] T. Zeller, P. Wild, S. Szymczak, M. Rotival, A. Schillert, R. Castagne, S. Maouche, et al. (2010) Genetics and beyond – the transcriptome of human monocytes and disease susceptibility. PloS One, 5, e10693.

